# Rapid neural encoding of the contrast between native and nonnative speech in the alpha band

**DOI:** 10.1101/2021.04.06.438632

**Authors:** Alessandro Tavano, Arne Nagels, Benjamin Gagl, R. Muralikrishnan

## Abstract

More than half of the world’s population is multilingual, yet it is not known how the human brain encodes the perception of native vs. nonnative speech. To find out, we asked German native speakers to detect the onset of native and nonnative (English and Turkish) vowels in a roving standard stimulation. Using EEG, we show that nonnativeness is robustly registered by an increase in phase coherence in the alpha band (8-12 Hz), beginning as early as ∼100 ms after stimulus onset and lasting more than 200 ms. The alpha band effect is speech-specific, successfully predicts the response speed advantage of nonnative speech, and grants ∼90% decoding accuracy in distinguishing native vs. nonnative speech irrespective of language familiarity. We propose alpha phase coherence as a candidate neural channel for the online resolution of the native-nonnative contrast in the adult brain.

**Significance Statement:** We show that the human brain takes only ∼ 100 ms to distinguish native vs. nonnative speech. This difference is neurally encoded in the alpha band in a reliable and specific way, supporting a processing model in which the mixture of top-down and bottom-up information represented in the speech signal is concurrently partitioned into different frequency channels.

## Introduction

Speech nativeness constitutes a uniquely robust life-time prior, capable of shaping strong social bonds or an inescapable diffidence (Grosjean, 2010), yet it is presently unknown which frequency channel encodes the native-nonnative contrast in speech perception in the adult brain. The theta channel (4-7 Hz) seems to play a central role in tracking rapid changes in speech envelope amplitude (Giraud and Poeppel, 2012). As the classification of native vs. nonnative speech relies on the spectrotemporal analysis of individual phonemes, the native-nonnative contrast could be encoded within the theta channel by changing its sensitivity to acoustic information. Indeed, the theta channel stands out as specialized for speech acoustics, while delta (1-3 Hz), alpha (8-12 Hz) and beta (13-30 Hz) neural frequency bands do not seem to carry acoustic information to a comparable degree (Teng and Poeppel, 2020). Alternatively, nonnativeness -conceived of as a violation of the nativeness prior -might activate a separate neural channel dedicated to top-down processing, working in parallel with the bottom-up acoustic processing within the theta band.

We asked young adult native German speakers (N = 27) to detect stimulus changes in an auditory stream of six vowels presented continuously every 800 ms, while their electroencephalographic (EEG) signal was recorded. Three vowels belonged to the German speech system (native), and three were nonnative (English and Turkish, Figure 1a). As speech is tracked in time, we used a roving standard stimulus design (Haenschel et al. 2005) to partial out the contribution of temporal orienting of attention from the native-nonnative contrast: each vowel appeared either three, six or nine consecutive times (standard = last element of a series), before changing into one of the other five (deviant = first element of the next series, Figure 1b).

**Fig 1.**
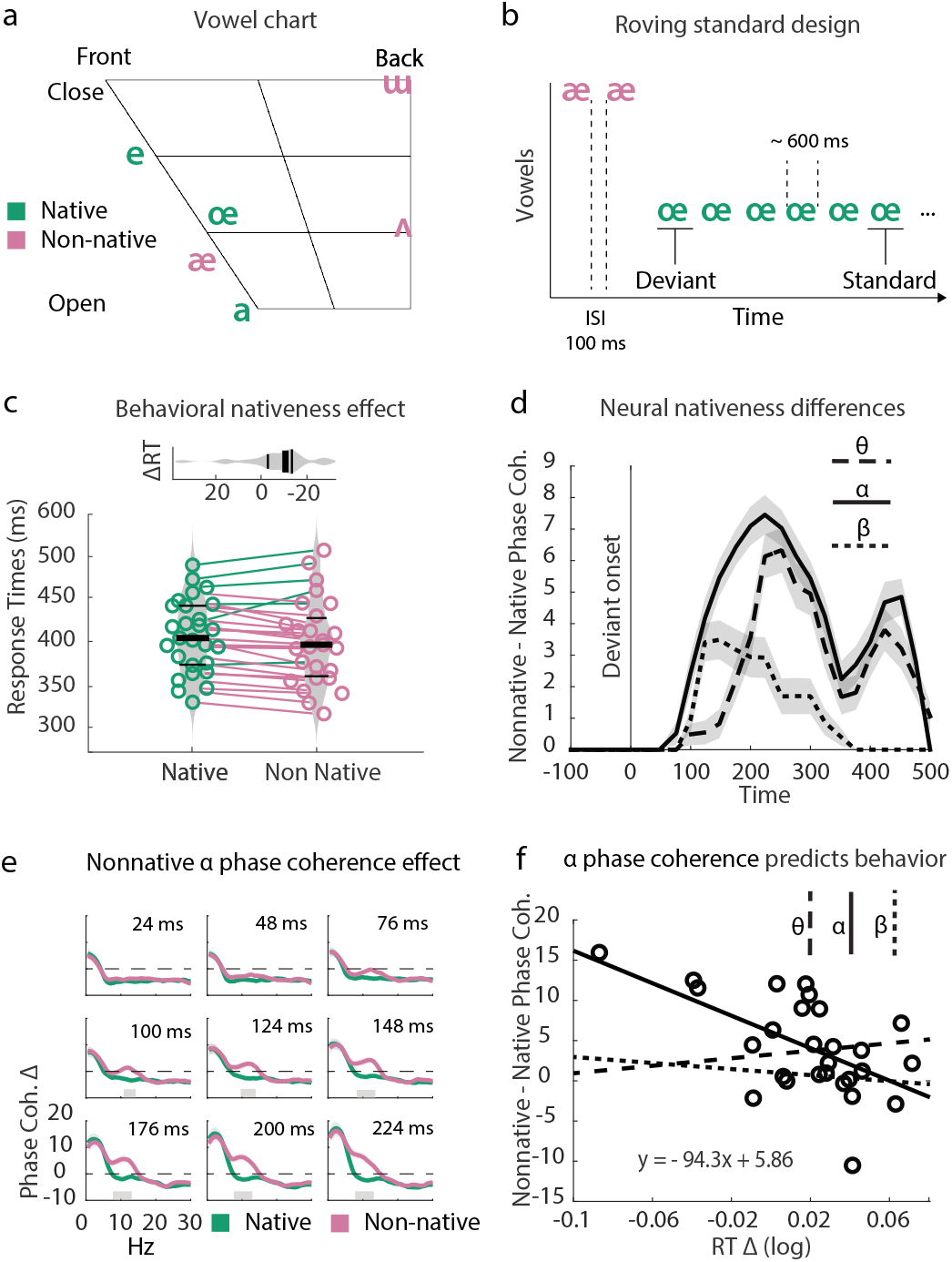
Alpha phase coherence predicts nativeness constrast. a. Vowel chart for the experimental vocalic stimuli http://www.ipachart.com/. Three were German vowels (green) and three non-German vowels (magenta), with two being English vowels (highly familiar, /æ/ and /Λ /), and one belonging to the Turkish language (/ш), and thus was relatively unfamiliar. Control stimuli were created by quilting the pristine ones in order to preserve spectral content (quilt segment = 10 ms, see Materials and Methods). All stimuli were root-mean-square equated for intensity. b. Stimuli were delivered using a roving standard paradigm, separately for vocalic and quilted controls. The first stimulus of each new sequence is termed Deviant, while the last is termed Standard, providing a perfect match in physical features. c. Response times show a mean advantage of nonnative stimuli of about 15 ms. Colored lines between groups indicate single-subject advantage in one or the other direction. The nativeness effect was independent of the orienting of attention in time (see text) d. A cluster-based permutation analysis highlighted significant differences in *θ, α* and *β* bands, plotted as the summation of all significant timefrequency points across all significant scalp electrodes. Shaded areas indicate ±Standard Error of the Mean (SEM). Phase coherence step-wise plots indicate that nonnative phase coherence increases post-stimulus specifically at alpha frequencies. Grey rectangles on the x axis indicate significant permutation clusters. f. By fitting the logarithm of the nativeness behavioral effect to the nativeness effects on phase coherence, the alpha band stands out as selectively significant. Circles represent individual participants for the alpha fit only. For the equation, x = RTΔ, y = *α*.

Since the roving standard paradigm allows perfect physical matching between standards and deviants, we contrasted deviant-minus-standard inter-trial phase coherence responses for native and nonnative vowels. Phase coherence has been shown to reliably reflects speech acoustic processing within the theta band (Teng and Poeppel, 2020), as well as deviancy detection and temporal attention (Tavano and Poeppel, 2019). Results show that the Nativeness factor is behaviorally independent from temporal attention to speech. Participants were faster at detecting the onset of nonnative deviant vowels, an advantage predicted by an increase in alpha band phase coherence. Importantly, neural decoding of alpha band phase coherence captured ∼90% of accuracy in distinguishing native vs. nonnative speech stimuli.

## Results

### Behavior

The temporal orienting of attention to stimuli benefited both accuracy and response speed in vowel-change detection regardless of nativeness: all *Fs*_(2,52)_ ≥13.69, all *ps* ≤ 0.001, partial *η*^2^ ⩾ 0.15. The Nativeness x Temporal attention interaction was not significant: *F* _(1,26)_ = 0.06, *p* = 0.94. Nativeness did not influence accuracy (*F* _(1,26)_ = 0.02, *p* = 0.88), but significantly modulated response speed: *F* _(1,26)_ = 8.35, *p* ¡ 0.01, partial *η*^2^ = 0.19. Participants were faster at detecting nonnative deviant vowels: mean response time for Native = 387.54 ms, SD = 40.35, Nonnative = 381.01, SD = 45.78. (Figure 1c). We conclude that the factors Temporal attention and Nativeness independently co-determine response speed in detecting vowel changes.

### A neural contrast between native and nonnative speech

We increased signal-to-noise ratio by separately averaging deviant and standard trials across the levels of the Temporal attention factor. Inter-trial phase coherence estimates for standard events were subtracted from those for deviant events to suppress the import of individual spectral differences. For all statistical purposes phase coherence differences were normalized using an arcsine transformation (Studebaker, 1985), however figures always display the original data. A cluster-based permutation test (Maris and Osstenvelt, 2007) between native and nonnative (Montecarlo resampling N = 2000), highlighted a significant post-stimulus Nativeness effect (p = 4.99^-04^, T = -5.42^+03^), which involved the Theta (4 -7.5 Hz) rhythm, the Alpha rhythm (8-12.5 Hz), and the Beta 1 rhythm (13-19.5 Hz). Figure 1d depicts band-wise mean phase coherence contrasts between nonnative and native trials across time, averaged across significant electrodes, and their dispersion: the alpha band clearly dominates the response from the earliest stages (peak phase coherence difference = 7.53%), but nonnative stimuli evoke also larger theta activity (peak phase coherence difference = 6.42%).

When broken down in time bins, the cluster-based significance shows an early increase in deviancy for nonnative trials at 100 ms centered on the alpha band (Figure 1e). This very value was confirmed by a conservative jackknife analysis of cluster onset (estimated as the earliest jackknife point, for values averaged across all electrodes, that is significant for all participants), indicating that from 100 ms a significant difference in the alpha band was found in all participants (significant cluster onset for theta band = 252 ms post-onset; the Beta 1 rhythm did not meet the strict onset threshold). Mean values calculated for ∼50 ms windows around bandwise peak maxima failed to significantly predict behavior: all *Fs*_(1,25)_ : ⩽ (2.87, all *ps ⩾* 0.10. However, when the peaks of the first derivative indicating maximal rate of change were used (∼50 ms window after non-parametric cluster onset: theta = 148 ms; alpha = 100 ms; beta = 100 ms), deviancy differences as reflected by phase coherence in the Alpha band explained a sizeable proportion of response time variance: R^2^ = 0.27, *F* _(1,25)_ = 9.35, *p* = 0.005. See Figure 1f: the linear equation weights (y = *α*; x = RTΔ) are based on original data plotted in Figure 1d. Theta and beta 1 signals did not predict behavior: all *Fs*_(1,25)_ ⩽ 0.85, all *ps ⩾*0.36.

### Decoding the native -nonnative speech contrast

We went on to classify native vs. nonnative stimuli from phase coherence differences in the alpha band. The set of nonnative stimuli is mixed relative to familiarity, as it includes English (highly familiar) and Turkish (less familiar) stimuli. To increase the precision of our findings and the experiment’s internal validity, we focused on the contrast between the highly familiar English stimuli /æ/ and / Λ/), and the two spectrally similar German ones (/œ/ and /a/). If familiarity plays a major role in nativeness perception, then the classifier should not perform above chance. We trained a multivariate pattern analysis (MVPA) classifier (linear discriminant analysis, LDA), with electrodes as a feature dimension to classify (decode) native (German) vs. nonnative (English) stimuli from the deviant-minus-standard differences in neural phase coherence. The performance of the classifier – scored by computing the nonparametric measure of the area under the curve (AUC) of the receiver-operating characteristic – was the highest (∼0.87 ROC) within the alpha band (∼8-10 Hz, Figure 2a, continuous line), and well above chance (0.5 ROC, classification resampling N = 1000).

**Fig 2.**
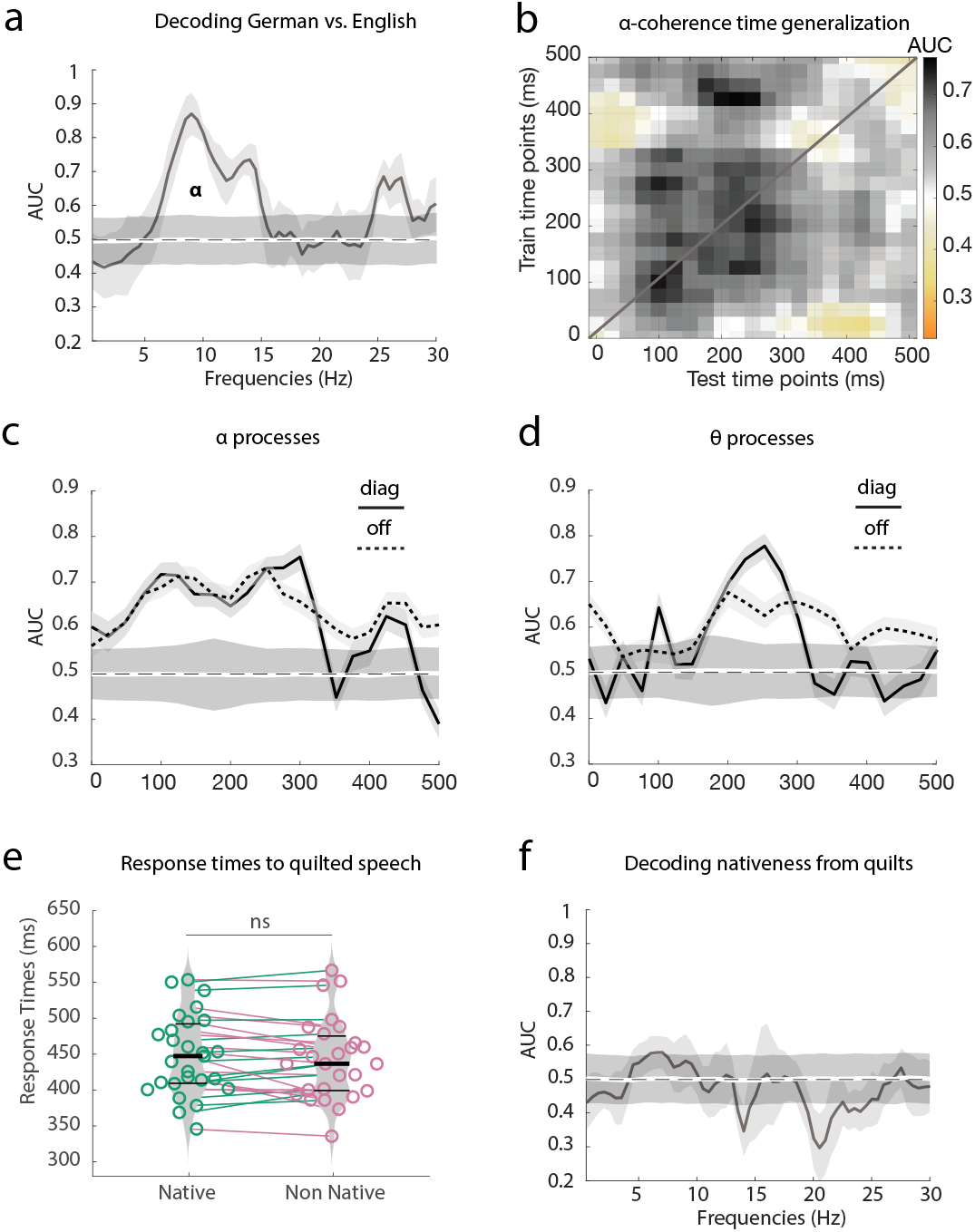
a. Nativeness was decoded from the contrast of deviant-minus-standard phase coherence responses, using an LDA classifier. Notice that the LDA classifier successfully distinguishes -with a large peak in the alpha range -German and English vowels that are very similar in spectrotemporal profile (see text). b. Time generalization shows a square profile for the alpha band starting between 100 and 300 ms, suggesting that the same neural process is maintained in time. c. The same data as in b., but this time distinguishing the diagonal and the (averaged) nondiagonal activations. d. For the theta band, no clear square pattern was detected. e. No significant diference in response times was found to the onset of native vs. nonnative quilted vowels (acoustic controls). f. Congruently, the LDA classifier failed in decoding German and English quilted vowels

We then generalized this approach over time (King and Dehaene, 2014) to test if an identified neural pattern at one time point would persist later on, indicating that the same type of information is shared across successive stages of stimulus processing.

Indeed, alpha phase coherence highlighted a square generalization pattern, with ∼ 0.7 ROC (Figure 2b). Figure 2c illustrates this point by plotting the activity on the diagonal, which assesses classifier performance at the time used for training, but also off diagonal activity, which assesses classifier performance on data from other time samples. Instead, theta decoding peaked on the diagonal at ∼ 250 ms (Figure 2d).

### The alpha effect is speech-specific

It is possible that the the alpha phase coherence effect reflect spectrotemporal novelty, rather than speech-specific differences. To test whether this is the case, we used quilting to subtract speechness quality from the original stimuli, while preserving their spectrotemporal content (Overath et al., 2015). A significant effect of Temporal attention was replicated for both accuracy and response times: *F* _(2,52)_ ⩾ 13.1, *p* ⩽ 0.001, partial *η*^2^ ≥0.25. However, Nativeness did not significantly modulate accuracy or response times, either as a main factor or in interaction with Temporal attention (all *Fs* ⩽ 2.31, all *ps ⩾* 0.10; response times are depicted in Figure 2e).

A cluster-based permutation test found a significant Nativeness effect (p = 0.0060, T = -1.338^+03^), centered on the theta band but reaching also the lower portion of the alpha band (8-10 Hz), suggesting larger deviancy values for quilted nonnative than for native stimuli: peak phase coherence difference for theta band was 6.08%, and for alpha band 3.19%. Theta-band cluster onset (assessed via jackknife) was similar for quilted and pristine vowels: pristine = 252 ms, quilted = 200 ms. However, for the alpha band the lag was substantially larger: pristine = 100 ms, quilted = 476 ms. Neither theta nor alpha cluster values for quilted vowels (average window 176-224 ms) predicted response times: all *Rs*^2^ ⩽ 0.06, all *Fs*_(1,25)_ ⩽ 1.81, all *ps ⩾*0.189. Congruently, the LDA classifier did not distinguish native from familiar nonnative stimuli, when strapped of their pristine speech quality (Figure 2f).

## Discussion

We used deviancy to access both acoustic and phonological features of speech stimuli (Phillips et al., 2000). By contrasting physically matched deviant-minus-standard phase coherence responses, we found that the native-nonnative contrast is encoded by an increase in alpha band phase coherence for nonnative speech. The neural pattern subtending such an effect becomes significant soon after stimulus onset, at ∼100 ms, and is maintained for about 200 ms thereafter, virtually connecting early sensory to later more cognitive processes. Importantly, the alpha phase coherence effect is not influenced by language familiarity, another important statistical prior (Fleming et al., 2014). This suggests that the internal tagging of vowel stimuli as native may depend on a form of automatic matching to a template, with mismatch triggering an increase in alpha phase coherence.

By quilting the experimental stimuli, we were able to test the effects of spectrotemporal deviancy in the absence of speechness: These too involved the theta band, and marginally the alpha band, but with a radically late onset time, and no relationship to behavior. We infer that: 1. The alpha effect is speech-specific; 2. The native-nonnative contrast for pristine vowels may build the alpha effect upon a pre-determined neural space, sensitive to spectrotemporal differences.

An increase in alpha phase synchrony has been shown to underlie increased communication between sensory, associative, sensorimotor and motor brain areas (Moore et al, 2008). Indeed, participants in our study were faster at detecting nonnative deviant vowels than native ones, and alpha phase coherence was a significant predictor of response speed.

We suggest that during on-line perception, separate neural channels concurrently run different operations: the theta channel may be predominantly deputy to acoustic feature analyses (Teng and Poeppel, 2020), while the alpha channel would react to violations of priors. Both processes in turn may depend on even faster processes -possibly subcortical in origin -declaring a mismatch to the nativeness prior based on fine spectrotemporal templates (Wartenburger et al., 2003; Parras et al., 2017). While this suggestion awaits future verification, the high sensitivity of alpha phase coherence to the native-nonnative speech contrast, and the fact that it is a robust signal, easy to sample, makes it readily suitable to study not only adults but also the maturational underpinnings of speech acquisition in infants and children immersed in bilingual settings.

## Acknowledgments

The authors gratefully acknowledge help from Miriam Riedinger with data collection, and the support of the Max Planck Society and the Johannes Gutenberg University, Mainz (DE).

## Materials and methods

### Participants

Participants were twenty-nine young adults (Age range: 18-36, mean = 24.9, 18 females), German native speakers, predominantly right-handed (adapted Oldfield test), self-reporting normal hearing, normal-to-corrected vision, and no medical history of CNS treatments, who were compensated for their participation. All experimental procedures were approved by the Ethics Committee of the Johannes Gutenberg University, and each participant signed a written informed consent to them. Two participants were excluded because of low accuracy scores (¡ 30%), and outlier response times (¿ 3 standard deviations, SD). We report on the results for the remaining 27 participants.

### Stimuli

Six vowels were extracted from ipachart.com/ under a Creative Commons licence, equated in length (700 ms), ramped 20 ms at either end, and matched for intensity using the root mean square. Three vowels belonged to the German speech system (native stimuli: /e/, /a/, and /œ/), while the remaining three did not (nonnative stimuli: /æ/, /Λ / and / w /. Nonspeech controls for each vowel were generated using a quilting-and-shuffling approach on the original stimuli (sound waves were divided into 360 segments of 85 data points each, corresponding to ∼1.9 ms; segments 24 to 300 were then randomly shuffled, while the initial and final speech fragments were kept intact (Overath et al., 2015).

Stimuli were repeated either three, six or nine instances and concatenated with an inter stimulus interval (ISS) of 100 ms (roving standard, Figure 1b; Haenschel et al, 2005). Each stimulus sequence contained 108 stimuli, and each participant received 20 stimulus sequences, 15 with clear vowels -native and nonnative -, and 5 with quilted vowels (acoustic controls). The same vowel/acoustic control could be repeated only if interspersed by three different vowels/acoustic controls. The first instance of a new stimulus is termed *Deviant stimulus*, while the last instance is termed *Standard stimulus*: this allows studying Standard and Deviant as purely statistical occurrences, abstracting from their matching physical features.

### Procedure

Participants (N = 27) were instructed to fixate a circle at the centre of the screen, listen attentively and press a button on a response box to the onset of each Deviant, regardless of type. Stimuli were delivered diotically at 75 dBs SPL via loudspeakers positioned at circa 1.2 meters from participants, and further attenuated using a fixed -20 dB SPL step. Stimulus sequences were created using custom scripts written in Matlab (R2015b, 64 bit, mathworks.com) and Presentation (Neurobehavioral systems, https://www.neurobs.com/). Sequence delivery was controlled by Presentation, running on a Windows XP computer.

## Data Recording

We used a passive electrode set with 24 Ag/Ag-Cl scalp electrodes (10-20 system, Easycap, http://easycap.de/), 4 external electrodes positioned at either eye-canthus, above and below the right eye to record horizontal and vertical eye movements (bipolar montage), and over left and right mastoid formations (N = 30 electrodes) to record electroencephalographic (EEG) activity (BrainAmp DC amplifier, brainproducts.com/) at a sampling rate of 1KHz. All impedances were kept below 10 kOhm. Data were recorded with an online reference to the left mastoid, and offline downsampled to 250 Hz and filtered 0.2 to 45 Hz (Kaiser window, filter orders 2267 and 93, transition bands 0.4 and 10 Hz; firfilt plugin for EEGLAB by Andreas Widmann). Electro-ocoular artefacts were detected and rejected by submitting the downsampled EEG data (high-passed filtered at 1 Hz) to an Independent Component Analysis using the Infomax algorithm. The resulting Independent Components (ICs) were tested using the SASICA toolbox for EEGLAB (Chaumon et al., 2015): ICs reflecting blinks/vertical eye movements and lateral eye movements were detected by means of a correlation threshold (0.7) with bipolar Vertical and Horizontal EOG channels, and found to be present in all participants (range: 1-2 ICs for vertical and 1-2 ICs for horizontal eye movements).

## Data Analysis

Behavioral detection accuracy was calculated for the number of detected deviant vowels, and together with response times for corrected detections was submitted to a 3×2 repeated measures Analysis of Variance (rmANOVA) with factors Temporal attention (Three levels: three, six, nine sounds in each standard series) and Nativeness (two levels: native vs. nonnative). Statistics were run on standardized data: Original data are used when reporting means and standard deviations, and for plotting. A logarithmic transformation was used when regressing behavior on neural data. Inter-trial phase coherence (ITPC) analysis was run using the Fieldtrip toolbox for Matlab (www.ru.nl/neuroimaging/fieldtrip), and custom Matlab scripts. The complex Fourier spectrum was obtained for epochs from 750 ms before onset to 1250 ms post-onset, then element-wise multiplied at each sensor with a Fourier transformed Morlet wavelet (Hanning taper) in the frequency domain (0.25 Hz resolution, from 0.25 to 40 Hz), in steps of 25 ms, and then inverse Fourier transformed. ITPC values (ranging from 0 to 1) were computed for each Nativeness level by amplitude-normalizing the Fourier output, summating the angles and normalising by length. ITPC is insensitive to differences in amplitudes between trials. For statistical purposes, phase coherence differences were turned into rationalized arcsine units (Studebaker, 1985).

Nativeness decoding was run on the phase coherence scores for each participant, separately for the theta and alpha bands. Three independent, numerically balanced subsets (“folds”) were created for each band from the scores of all participants (“samples”), and an iterative process was repeated independently for each fold, while the other two were used as training set to train an LDA (Linear Discriminant Analysis) classifier (with electrodes as features). The resulting classifiers were tested by predicting whether a trial label was native or nonnative. Accuracy was quantified using the area under the curve of the receiver-operating characteristic (ROC AUC) This analysis was repeated three times, with each trial subset serving as testing set once. A Montecarlo resampling approach (1000 repetitions) was used to obtain 95% confidence intervals, by shuffling the order of the trial labels and replicating the LDA classification.

## Data and code accessibility

Data and code used for statistical analysis, figures and detailed analysis protocols at the following address: osf.io/2haxk/?view_only=b1e8c5af6b074c41a22c2d4c5366b291.

